# Mathematical models of protease-based enzymatic biosensors

**DOI:** 10.1101/695320

**Authors:** Deepak K. Agrawal, Sagar D. Khare, Eduardo D Sontag

## Abstract

An important goal of synthetic biology is to build biosensors and circuits with well-defined input-output relationships that operate at speeds found in natural biological systems. However, for molecular computation, most commonly used genetic circuit elements typically involve several steps from input detection to output signal production: transcription, translation, and post-translational modifications. These multiple steps together require up to several hours to respond to a single stimulus, and this limits the overall speed and complexity of genetic circuits. To address this gap, molecular frameworks that rely exclusively on post-translational steps to realize reaction networks that can process inputs at a timescale of seconds to minutes have been proposed. Here, we build mathematical models of fast biosensors capable of producing Boolean logic functionality. We employ protease-based chemical and light-induced switches, investigate their operation, and provide selection guidelines for their use as on-off switches. We then use these switches as elementary blocks, developing models for biosensors that can perform OR and XOR Boolean logic computation while using reaction conditions as tuning parameters. We use sensitivity analysis to determine the time-dependent sensitivity of the output to proteolytic and protein-protein binding reaction parameters. These fast protease-based biosensors can be used to implement complex molecular circuits with a capability of processing multiple inputs controllably and algorithmically. Our framework for evaluating and optimizing circuit performance can be applied to other molecular logic circuits.

## Introduction

Cellular signaling networks perform efficient computation in a complex environment by sensing and processing a multitude of chemical and physical signals into various responses that play a central role in cell metabolism and function. This biological computation has inspired the creation of synthetic genetic networks that follow defined input-output characteristics to generate a diversity of response outputs in response to a set of inputs as a proof-of-principle for biological computation.^1–3^ These genetic networks require transcription and translation reactions (after input sensing) to produce proteins. The period of performance of a single input-output layer is several hours. These output proteins (e.g. transcription factors) can be used in turn, to control the expression of other genes in cascade or multi-layer network topologies. However, the long (∼hours) time-scales involved preclude efficient construction of multi-layer networks. For biosensing applications where rapid sensing and response are required,^4,5^ genetic networks are, thus, of limited value. Moreover, heterogeneity of the intra-cellular environment and crosstalk between synthetic and endogenous components limit not only the speed but also the overall robustness of circuits based on transcription.^3^ An alternative biomolecular component library that can potentially operate at a much faster timescale compared to transcription-based genetic networks may help realize complex circuitry for situations, such as real-time detection, where a fast and precise response is required.

Recently, an alternative paradigm has been developed to realize fast biomolecular computations. This approach uses recombinantly expressed synthetic proteins fused to enzyme fragments.^6^ Unlike transcription-translation reactions, enzyme catalysis occurs at a timescale of seconds to minutes, and thus protein-based enzymatic biochemical circuits operate at much faster timescales than the transcription-based circuits ^7^ (Fig. 1a and b). Moreover, one enzyme molecule can process multiple substrate molecules by virtue of catalytic turnover; in contrast, only stoichiometric binding is possible with transcription factors. Therefore, large signal amplification is expected in enzyme-based circuits. These features led to the development of several enzyme-based reaction networks to implement analog and digital logic functions;^8^ yet, there are few generalizable approaches that can be robustly tailored to implement biosensors with a capability of processing multiple inputs at tunable speeds and controllable sensitivity.^9–11^ Many enzymatic circuits are built ad-hoc, using specific substrates/products of particular enzymes as input/output signals.^8,9,12,13^ However, recent advances in the construction of entirely bio-orthogonal, post-translationally responsive and controllable protease-based systems have enabled the development of modular components that can potentially be used as building blocks for generating fast and controllable biomolecular circuits both *in vitro* and *in vivo*.^3^

**Figure 1:**
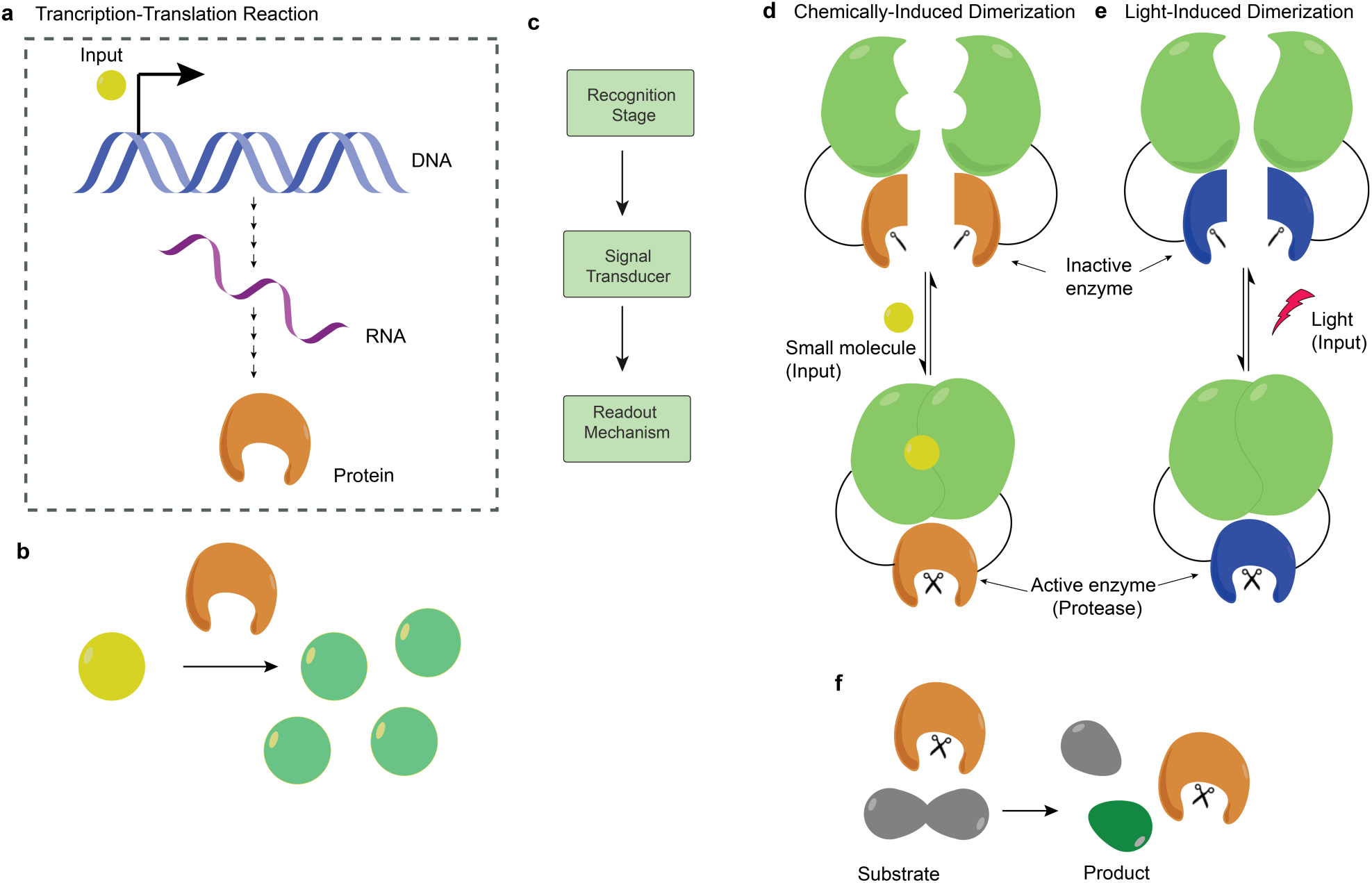
Enzyme-based reaction networks. **(a)** Genetic regulatory circuits typically use gene expression, which requires transcription (converting DNA to RNA), and translation (converting RNA to protein) reactions. The requirement for these steps leads to long timescales for circuit operation (hours-days) **(b)** In contrast, enzyme catalysis, where a protein enzyme acts as a catalyst for converting substrate molecule into a product, occurs on seconds to minutes timescales. **(c)** Block diagram representation of the biosensor. **(d-e)** Concept of induced molecular dimerization. In this scheme, two inactive fragments of a protease are attached to two different protein molecules, which cooperatively undergo dimerization in the presence of **(d)** chemical or **(e)** optical signals to induce folding of the attached enzyme fragments and restore its activity. **(f)** Quenched fluorescent protein system in which only an active protease can cleave the substrate, leading to an increased fluorescence signal.

In this paper, we present a bottom-up design framework for the rapid detection and response to chosen chemical and optical inputs using protease-based logic circuits. We first design two elementary circuits (called switches) based respectively upon chemical and light-induced dimerization mechanisms and capable of producing the same type of output in response to different kinds of inputs. We evaluate their responses for a broad range of reaction conditions and provide screening criteria to optimize their performance. We then use these switches to design biosensors, which can process two different types of inputs simultaneously and produce an output that follows either OR or XOR Boolean logic functionality. We develop comprehensive ordinary differential equation (ODE) models for these biosensors and analyze their dynamic response through numerical analysis at a realistic set of reaction parameters. We further conduct sensitivity analysis to determine the influence of reaction parameters on the output dynamics over time. Our results indicate that a variety of digital signal processing functions can be implemented by using these switches as elementary blocks. Additionally, our approach illustrates several quantitative methods useful for assessing the performance of biomolecular logic gates.

## Results

### Designing biosensor components

Our overall goal is to design biosensors that can process two different types of inputs and rapidly produce a well-defined output signal. The output should be tunable to meet performance specifications that govern the overall dynamics of the system. For example, the biosensor should be able to differentiate between cases where both inputs are absent and the cases where either or both inputs are present (an OR gate). The OR gate design should be extendable to achieve an XOR Boolean logic function, which allows to differentiate between cases where both inputs are either absent or present and cases where only one input is present at a time.

As a starting point, we choose to divide the operation of the biosensor into three modules (Fig. 1c); recognition stage, signal transducer, and read-out mechanism. A recognition stage detects the presence of a particular input, which in our design can be chemical or physical. A signal transducer then converts the detected signal resulting from the interaction between the input and the recognition stage into a measurable molecular activity. In the final stage, the readout mechanism allows generating and reading an optical signal corresponding to the output.

To process chemical and physical input signals via the recognition stage, we used chemically and light induced dimerization mechanisms respectively. In chemically induced dimerization (CID), a small molecule or a dimerizer acts to bring two proteins or protein fragments together, leading to an increase in the effective concentration of dimerized complex. ^14,15^ CID mechanisms have been widely used to rapidly manipulate molecular activities in cells from a variety of species, including *E. coli*. Similar to the technique of CID, light-induced dimerization (LID) exploits a pair of specialized protein domains that can be driven into a high-affinity binding state by illumination with a specific wavelength of light.^16^ Light-induced dimers are especially useful because they can be turned on and off with high spatial and temporal resolution in living systems, allowing for control of protein localization and, in our application, enzyme activity. Several LIDs are currently available and have been used to control signaling pathways in living cells.^17–20^

For signal transduction, split proteases can be fused to the pair of proteins used by CID and LID reactions.^21,22^ Each split protease is inactive on its own. In the presence of the input, the pair of proteins come together to form a dimer. While this happens, the split proteases come in close proximity, which allows them to reconstitute and thereby restore protease activity (Fig. 1d and e). The signal transduction along with the recognition stage allows converting a chemical or physical input into a sufficient concentration of the active protease. Finally, for a read-out signal, we use a green fluorescent protein (GFP) which is fused to a quencher protein. The reconstituted protease can cleave the link between GFP and the quencher protein, which causes an increase in the measured fluorescence signal (Fig. 1f). In the absence of the active protease, the fluorescent signal is negligible as the quencher is in close proximity to the fluorescent protein. ^23^ Signal amplification occurs when the transducer enzymatically converts the protease concentration into GFP output. Our approach is to connect the recognition stage, signal transducer and the read-out mechanism to build mathematical models for the elementary blocks, which can produce a GFP output in response to either chemical or optical signal. These blocks can then be used to design biosensors capable of processing two inputs simultaneously.

### Modeling CID switch

To process a chemical signal, we start by developing a model for a CID switch. To be realistic in our approach, we considered a switch that uses FK506 binding protein (FKBP) and the FKBP rapamycin binding protein (FRB) as the two protein fragments. These two protein fragments can form a dimer in the presence of rapamycin. ^24–26^ By fusing a fragment of an inactive protease with each protein, that is otherwise unfolded but folds upon enhanced proximity with one another, rapamycin-dependent enzyme activity can be observed. The high affinity of the ternary complex means that small concentrations of rapamycin can be used to trigger enzyme folding, and the entire action can be induced on a timescale of seconds. A previous report of such a CID-split protease fusion shows that Tobacco Etch Virus (TEV) protease activity can be robustly reconstituted in response to rapamycin addition by fusing fragments of TEV protease to FKBP and FRB proteins.^26^ For the read-out signal, we considered a recombinant intramolecular FRET construct consisting of GFP and resonance energy-accepting chromoprotein (REACh), fused by a recognition peptide sequence for TEV protease. TEV can cleave this sequence, causing the GFP output to increase.^23^ The concentration of the GFP output, therefore, depends on the concentration of the input (rapamycin).

In the presence of input, an interaction between FKPB (*A*) and FRB (*B*) proteins leads to the formation of TEV protease (*E*), which then cleaves the GFP-REACh substrate (*S*) to produce the GFP output *P* (Fig. 2a). The molecular interactions involved in the formation of the FKPB-rapamycin-FRB ternary complex have been investigated earlier and demonstrated to have two different pathways through which the ternary complex can form (Fig. 2a and b). ^25^ Moreover, split TEV protease fragments can interact with low affinity to generate (at low levels) the active enzyme even in the absence of rapamycin (*R*).^22^ We refer to this rapamycin-independent association of the components as a “leak reaction”. To understand the operation of the CID switch, we model its kinetics using an ODE model (see Supplementary Note S1) at a feasible set of reactions parameters (Table 1). For simplicity, hereon FKBP, FRB, rapamycin, TEV, GFP-REACh and GFP are denoted as *A, B, R, E, S* and *P* respectively. The subscripts 0 corresponds to the initial concentration of the respective species except for *P*. We determined the switch response in the absence and presence of *R*, which are denoted as 0 and 1 respectively. We observed almost the same output response for input 0 as for input 1 (Fig. 2c). The lack of difference is likely due to the leak reaction between *A* and *B*, which takes place even in the absence of *R* and leads to the production of *E*^′^, which then reacts to *S* to produce *P* (Fig. 2b).

**Table 1:** Model parameters for each reaction network. In this study, 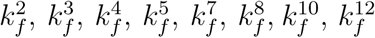 were same as 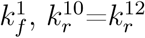, and 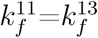

| Parameters | Values | Units |
| --- | --- | --- |
| $k_f^1$ | $1 \times 10^6$ | $\text{M}^{-1}\text{s}^{-1}$ |
| $k_r^1$ | $2 \times 10^{-4}$ | $\text{s}^{-1}$ |
| $k_r^2$ | 26 | $\text{s}^{-1}$ |
| $k_r^3$ | 12 | $\text{s}^{-1}$ |
| $k_r^4$ | $2 \times 10^{-3}$ | $\text{s}^{-1}$ |
| $k_r^5$ | 74 | $\text{s}^{-1}$ |
| $k_f^6$ | 0.02 | $\text{s}^{-1}$ |
| $k_r^7$ | 20 | $\text{s}^{-1}$ |
| $k_r^8$ | 0.8 | $\text{s}^{-1}$ |
| $k_r^9$ | 47 | $\text{s}^{-1}$ |
| $k_r^{10}$ | 0.01 | $\text{s}^{-1}$ |
| $k_f^{11}$ | 1 | $\text{s}^{-1}$ |

**Figure 2:**
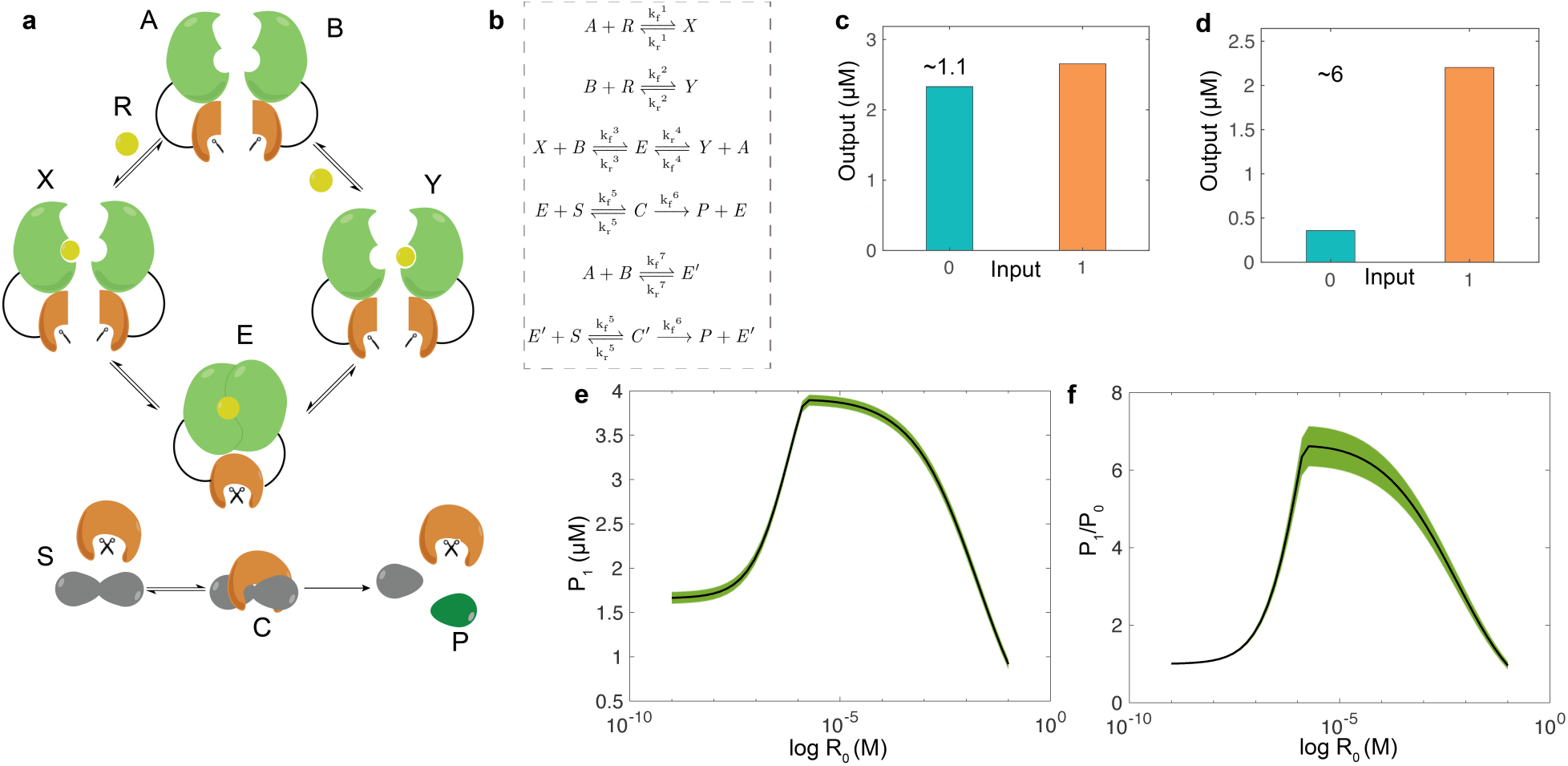
Small molecule induced switch. **(a)** Schematic diagram represents a switch that uses the chemically-induced dimerization (CID) mechanism. In the presence of rapamycin molecule (*R*), FKBP (*A*) and FRB (*B*) proteins form a complex, which leads to restore the activity of split protease *E*. The protease *E* then cleaves a fluorophore substrate (*S*) to produce green fluorescent protein output (*P*). **(b)** The corresponding chemical reaction equations for the CID switch. We derived an ODE model (see SI Note S1) from the chemical reactions and used the model to simulate the response of the CID switch with parameters shown in Table 1. **(c-d)** Simulated response of the CID switch at the initial concentrations of **(c)** *A*_0_=*B*_0_=5 *µ*M, and *R*_0_=0.5 *µ*M; **(d)** *A*_0_=*B*_0_=1.44 *µ*M, and *R*_0_=3.01 *µ*M. Here, 0 and 1 represent the absence (*R*_0_=0 *µ*M) and presence of the input signal (*R*), *S*_0_ was 5 *µ*M while the rest of the molecular species were initially set to 0 *µ*M. (e-f) Performance evaluation of the CID switch at different values of *R*_0_ while keeping *A*_0_ and *B*_0_ fixed at 1.44 *µ*M each. For each concentration of *R*_0_, 1,000 simulations were conducted where we randomly sampled a set of parameter values from a uniform distribution (See Methods). Averaged metrics of *P*_1_ and the *P*_1_/*P*_0_ are shown in **(e)** and **(f)** respectively. All the values of *P* were determined at 30 min. The error bars are shown in the shaded region and were determined using the standard error of the mean.

### Changing the reaction conditions allow to improve the response of the CID switch

From a switch, typically, a high output (*P*_1_) is desired when the input is 1 (presence of *R*) compared to a low output (*P*_0_) when the input is 0 (absence of *R*). This requires defining a specific output range for each input condition for efficiently characterizing the switch operation as off (*P*_0_) or on (*P*_1_). Therefore, we set the following prerequisite conditions; a high output *P*_1_ should be more than 1 *µ*M, and a low output *P*_0_ should be less than 0.5 *µ*M, so that *P*_1_/*P*_0_ > 2. To achieve a response from the CID switch that meets our specification, initial concentrations of *A, B* and *R* were optimized (See Methods), and a desired response was achieved at reduced concentrations of *A*_0_, and *B*_0_, and an increased concentration of *R*_0_ compared to the earlier case (Fig. 2d) without changing the reaction parameters. At reduced initial concentrations of *A* and *B*, the total amount of *E* and *E*′ also reduces, but an increase in *R*_0_ increases only the amount of *E* (See Fig S1). Notably, the output of the transduction stage (*E*) takes only a few seconds to reach a steady-state (see Fig. S2). It should be noted that *P*_1_ and *P*_0_ values can be different from the ones we selected here as long as they can be measured accurately and it is easy to differentiate *P*_1_ from *P*_0_.

To understand the CID switch operation comprehensively, we evaluated its performance at a wide range of reaction parameter values, considering the absolute value of *P*_1_ and the *P*_1_/*P*_0_ ratio as metrics. To ensure that our results were not specific to the particular parameter value, we performed 1000 simulations where each parameter was randomly sampled from a uniform distribution from a bounded interval (See Methods) at the optimized initial concentrations of *A* and *B*. For simplicity, only the average metrics of *P*_1_ and the *P*_1_/*P*_0_ are shown in Fig. 2e and f respectively. The result obtained at a different value of *A*_0_ and *B*_0_ is shown in Fig. S3.

The response curves reveal that the switch performed best in terms of much higher values of *P*_1_ and the *P*_1_/*P*_0_ at a specific concentration of *R*_0_ (Fig. 2e and f). This is because the recognition stage of the CID switch has two possible reaction pathways through which *E* can form. An increase in *R*_0_, increases the amount of *E* as the interaction between *A* and *R* is much stronger than the interaction between *B* and *R* (See Table 1). However, after a critical point, increase in *R*_0_, led to a higher interaction between *B* and *R* and because of the fact that less *B* is available to bind to *X* in order to form *E* (see Fig. S4). Moreover, at the lower concentrations of *R*_0_, *P* is much higher than at the higher concentrations of *R*_0_ because at lower values of *R*_0_, *A* and *B* are freely available to produce *E* (product of the leak reaction) than at the higher values of *R*_0_ where *A* and *B* are sequestered by *R*, and forms *X* and *Y* (see Fig. S5). Finally, following the prerequisite conditions (*P*_1_ >1 *µ*M and the *P*_1_/*P*_0_ >2), we found that the minimum detectable concentration of rapamycin is 0.2 *µ*M and the range of detection is 15.3 mM. By changing this specification (*P*_1_ >0.5 *µ*M and the *P*_1_/*P*_0_ >2), it is possible to reduce the minimum detectable concentration to 2.1 nM, and simultaneously increase the range of detection to 32.4 mM (see Fig. S3).

### Modeling and improving the LID switch response

Our goal is to design biosensors that can process two different kinds of inputs simultaneously. We, therefore, sought to design a LID switch to detect a physical signal such as optical light. To build a realistic model of a LID switch, we considered iLID micro and SSPB, which form a dimer in the presence of blue light.^17^ This reaction has been investigated earlier, and a ∼50-fold increase in the concentration of the dimer was reported in the presence of blue light.^17^ Similar to the CID switch, by fusing fragments of the inactive split protease with each protein, it should be possible to robustly recover the active protease in response to a physical light signal. For simplicity, we assume that this protease, which is denoted as *F*, is orthogonal to the one used by the CID switch and forms ∼50 × more in the presence of input compared to when it is absent. Similar to *E, F* cleaves the substrate *S* to produce a GFP output *P* (Fig. 3a and b). Hereon, iLID and SSPB proteins are denoted as *U* and *V* respectively.

**Figure 3:**
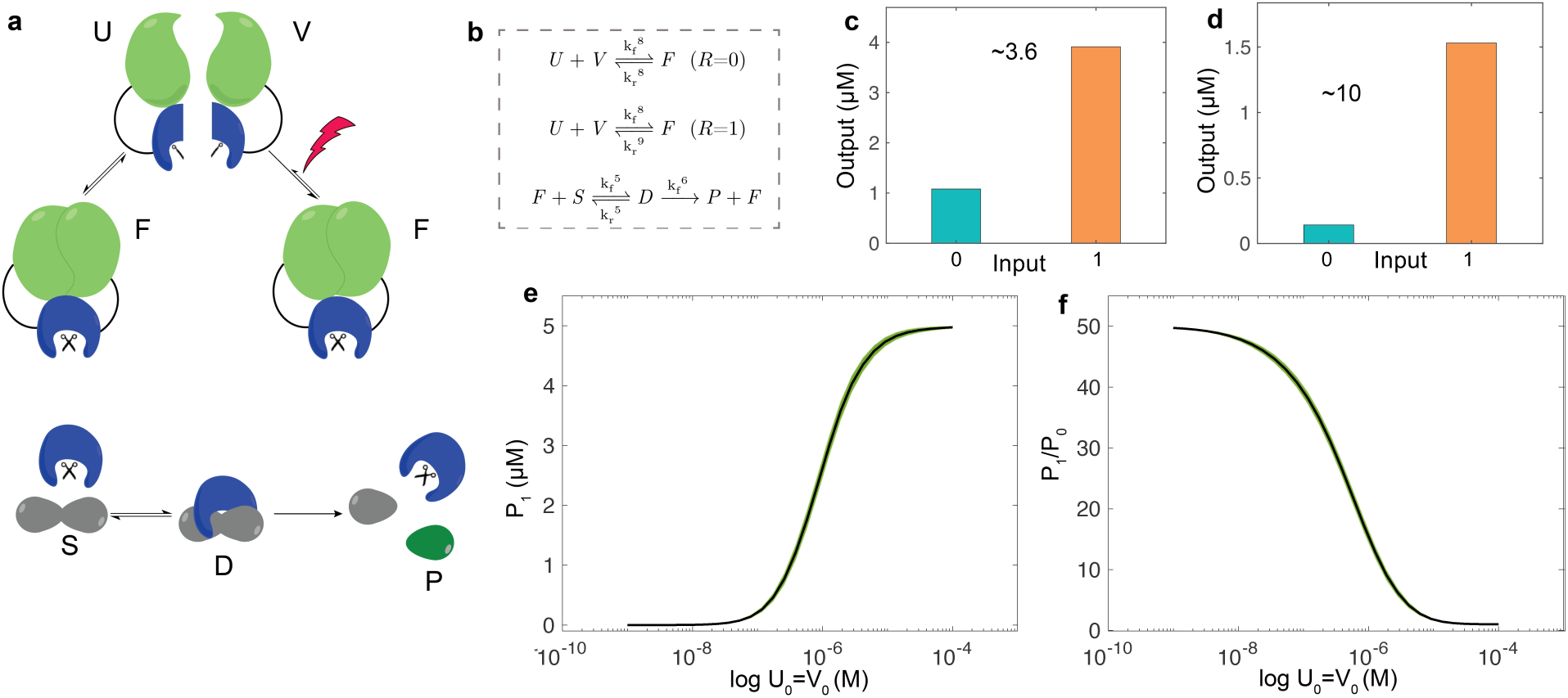
Light-induced switch. **(a)** Schematic diagram represents a switch that uses the light-induced dimerization (LID) mechanism. In the presence of an optical signal (blue light), iLID micro (*U*) and SSPB (*V*) proteins form a complex, which leads to the formation of a different version TEV protease, denoted as *F*. **(b)** The corresponding chemical reaction equations for the LID switch. We derived an ODE model (see SI Note S2) from the chemical reaction equations and used the model to simulate the response of the LID switch with parameters shown in Table 1. **(c-d)** Simulated response of the LID switch at the initial concentrations of **(c)** *U*_0_=*V*_0_=5 *µ*M; **(d)** *U*_0_=*V*_0_=1.59 *µ*M. Here, 0 and 1 represent the absence and presence of the input signal (*R*), *S*_0_ was 5 *µ*M while the rest of the molecular species were initially set to 0 *µ*M. (e-f) Performance evaluation of the switch at different values of *U*_0_ and *V*_0_ (*U*_0_=*V*_0_). For each initial concentration value, 1,000 simulations were conducted where we randomly sampled a set of parameter values from a uniform distribution (See Methods). Averaged metrics of *P*_1_ and the *P*_1_/*P*_0_ are shown in **(e)** and **(f)** respectively. All the values of *P* were determined at 30 min. The error bars are shown in the shaded region and were determined using the standard error of the mean.

We used an ODE model (see Supplementary Note S2) to determine the response of the LID switch at typical initial concentration values of *U* and *V* (denoted as *U*_0_ and *V*_0_), and found that *P*_0_ was more than 1 *µ*M (Fig. 3c). As per our prerequisite conditions, a low *P*_0_ value should be less than 0.5 *µ*M (*P*_1_ > 1 *µ*M and *P*_0_ *<* 0.5 *µ*M). A high *P*_0_ value was observed because of the undesired interaction between *U* and *V* in the absence of the input, that led to form substantial amounts of *F* (see Fig. S6). We then found reduced *U*_0_ and *V*_0_ values through optimization at which the switch response agreed with our specification (Fig. 3d).

To understand these results further, we determined the response curve for the LID switch as a function of *U*_0_ and *V*_0_ (*U*_0_=*V*_0_) at different random sets of reaction parameters (see Method). The mean performances are shown in Fig. 3e and f. Unlike the CID switch, the LID switch demonstrated completely different input-output characteristics (Fig. 2 and Fig. 3). For the LID switch, we observed that *P*_1_ was reduced with reduction in *U*_0_ and *V*_0_, but the *P*_1_/*P*_0_ ratio increased (within a bound). This means that lowering *U*_0_ and *V*_0_ led to a substantial reduction in *P*_0_ compared to *P*_1_. The binding affinity between *U* and *V* can be quantified using the dissociation constant (*k*_*d*_). The reported *k*_*d*_ value of *U* (iLID) and *V* (SSBP) is 47 *µ*M in the absence of light compared to 0.8 *µ*M in the presence of light.^17^ Typically, a smaller value of *k*_*d*_ suggests a high affinity between the two species and so by definition, if (*U*_0_=*V*_0_) *< k*_*d*_, the reaction is unlikely to form any *F*. Therefore, at reduced values of *U*_0_ and *V*_0_, the leak reaction produced negligible amounts of *F* (see Fig. S7).

### Designing a biosensor with Boolean OR gate functionality

One remarkable property of the CID and LID switches is the potential to network them together to make more complex circuits that can process versatile inputs. As a model system, we aim to design a protease-based biosensor that can mimic a Boolean OR gate functionality. A typical OR gate conventional symbol and a truth table are shown in Fig. 4a and b respectively. It has two inputs and one output, and can produce a high output only when either or both inputs are high. Such a response can, therefore, be used to detect the presence of either or both the inputs simultaneously. To design an OR gate based biosensor, we combined CID and LID switches where rapamycin and light inputs control the production rate of *E* and *F* proteases respectively. These proteases can cleave the same substrate *S* to provide a common output *P*. The two inputs, which are rapamycin and light, therefore, control the production of output *P* (Fig. 4c).

**Figure 4:**
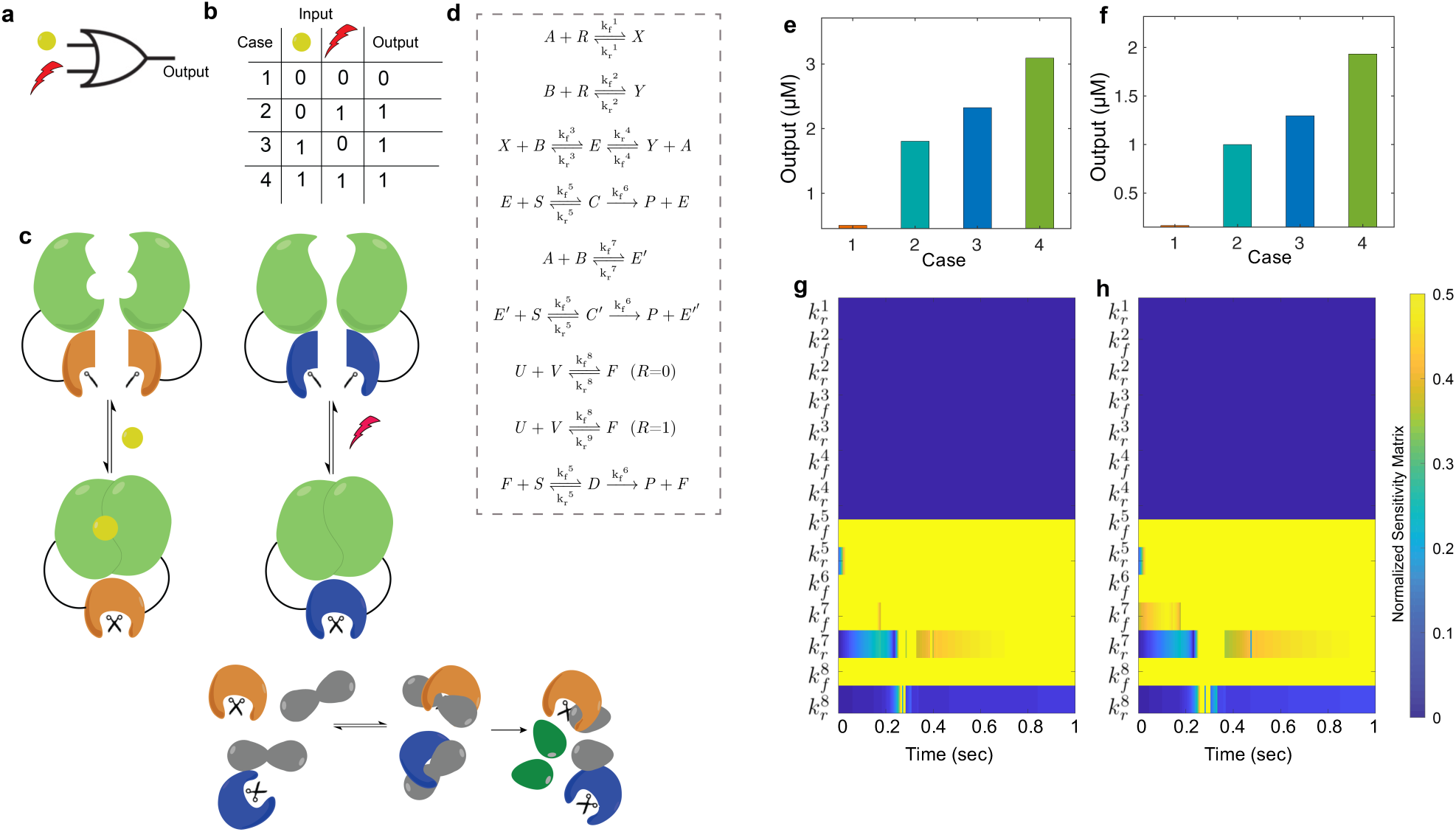
Protease based Boolean OR gate. **(a)** Conventional symbol and **(b)** a truth table of a Boolean OR logic gate. **(c)** Design of an OR gate that uses CID and LID switches, and **(b)** the corresponding chemical reactions. The OR gate has two inputs, rapamycin and light, that restore the activity of two split proteases *E* and *F* respectively. A high output in terms of the amount of GFP (*P*) results in when either or both the inputs are present demonstrating the operation of an OR gate. We model the kinetics of the OR gate using an ODE model (see SI Note S3) with parameters shown in Table 1. (e-f) Simulated response of the OR gate at **(e)** unoptimized (*A*_0_=*B*_0_=1.44 *µ*M, *R*_0_=3.01 *µ*M, and *U*_0_=*V*_0_=1.59 *µ*M) and **(f)** optimized (*A*_0_=*B*_0_=0.72 *µ*M, *R*_0_=1.67 *µ*M, and *U*_0_=*V*_0_=1 *µ*M) conditions and the corresponding results of the sensitivity analysis in **(g)** and **(h)** respectively for case 1 (both the inputs are absent). Normalized sensitivity matrix is shown with respect to the output (*P*). Here, yellow and blue correspond to the most sensitive and least sensitive values respectively. All the values of *P* were determined at 30 min. For cases 1 and 2, *R*_0_=0 *µ*M.

We next sought to model the kinetics of the chemical reaction network for this biosensor (Fig. 4d) for four different cases (Fig. 4b). In case 1, both the inputs are absent, while in cases 2 and 3 only the light signal or rapamycin is present respectively and in case 4, both the inputs are present. An OR gate operation requires a high output in cases 2, 3 and 4, and a low output in case 1 (Fig. 4b). We follow the same specifications as used earlier to categorize the output as either high or low (*P*_1_ > 1 *µ*M and *P*_0_ *<* 0.5 *µ*M). Using an ODE model (see Supplementary Note S3), we simulated the OR gate based biosensor response at the reaction conditions that were used to achieve the desired response from the CID and LID switches separately. Even though high responses were observed for cases 2, 3 and 4, in case 1 the output was more than 0.5 *µ*M, which contradicts our specification. We, therefore, sought to reduce the output activity when neither input is present while simultaneously maintaining a high output when either or both the inputs are present. To achieve this, we optimized the initial concentrations of *A, B, R, U* and *V*, and found an OR gate response that met our screening criteria without changing any reaction parameters (Fig. 4f).

The operation of an interconnecting reaction network with several species and reaction parameters can be challenging to understand, especially, in the presence of undesired interactions such as the leak reactions. To carry-out our optimization, it is required to determine how the reaction parameters govern the output dynamics at different reaction conditions for the same input combinations. For this purpose, we used sensitivity analysis to get an insight into how each model parameter affects the dynamics of the system. We, therefore, calculated the time-dependent sensitivity coefficient matrix to measure how sensitive the output is with respect to each parameter over time (see Methods). ^27,28^ The output sensitivity for the input combination in case 1 is shown in Fig. 4g and h before and after optimization respectively. The coefficients with a high value indicate that variations in the associated parameter cause a significant change in the output dynamics. Note that the coefficients of the normalized sensitivity matrix depend on time.

In Fig. 4g and h, high sensitivity of 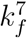 and 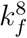 suggest that the leak reactions contributed to form the proteases (*E*^′^ and *F*) which then led to produce a high output value even in the absence of the inputs. At the optimized condition, a reduced sensitivity of 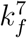 and an increased sensitivity of 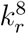 suggest a reduction in the amount of *E*′ and *F*, which resulted in lowering the output value (Fig. 4f and d). The output sensitivity for the rest of the three cases is shown in Fig. S8.

### Extending the OR gate design to achieve Boolean XOR gate functionality

A biosensor capable of processing a complex computation requires a complex circuit with a capability where different logic gates can read the same combination of inputs to produce an entirely different logical functionality. Our approach is advantageous over others^8^ in the sense that instead of designing a completely new reaction network for each logic gate, CID and LID switches can be considered as elementary blocks to design new logic gates. To demonstrate this capability of our approach, we aim to design a biosensor that can mimic an XOR gate functionality. The conventional symbol and a truth table of an XOR gate are shown in Fig. 5a and b respectively. Unlike the OR gate, which produces a high output when either or both the inputs are high (Fig. 4b), an XOR gate provides a high output when only one input is high (Fig. 5b).

**Figure 5:**
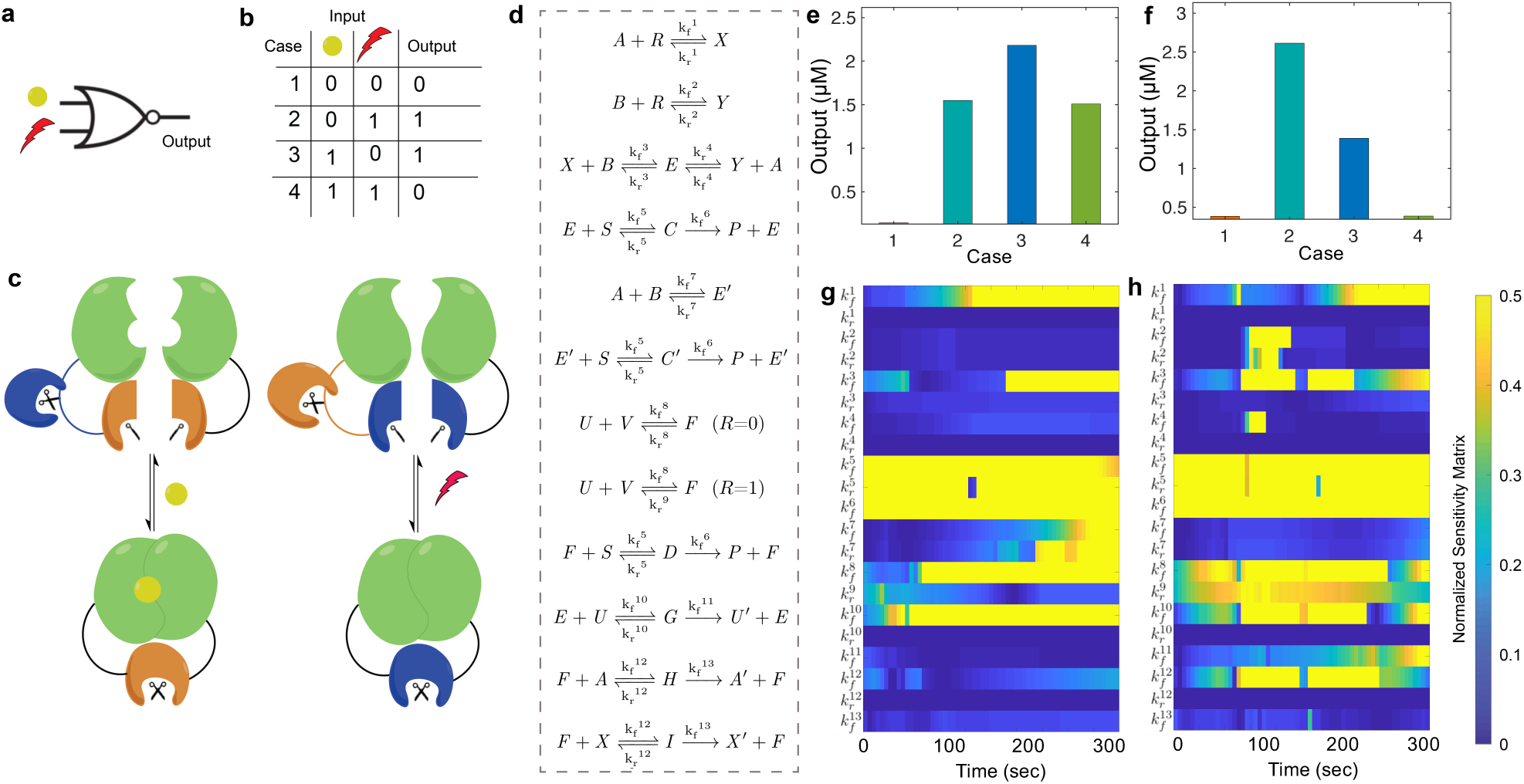
Protease based Boolean XOR gate. **(a)** Conventional symbol and **(b)** a truth table of a Boolean XOR logic gate. **(c)** Design of an XOR gate, which uses the CID and LID switches, and **(d)** the corresponding chemical reactions. Similar to the OR gate, the XOR gate has two inputs, rapamycin and light, that restore the activity of two split proteases *E* and *F* respectively. However, each of these two proteases can also inactivate the other enzyme by degrading the protein fragment orthogonally. These reactions limit the production of the two proteases when both the inputs are present, resulting in a low output. We model the kinetics of the XOR gate using an ODE model (see SI Note S4) with parameters shown in Table 1. Simulated response of the XOR gate at **(e)** unoptimized (*A*_0_=*B*_0_=1.44*µ*M, *R*_0_=3.01 *µ*M, and *U*_0_=*V*_0_=1.59 *µ*M) and **(f)** optimized (*A*_0_=1.22 *µ*M, *B*_0_=0.88 *µ*M, *R*_0_=0.94 *µ*M, and *U*_0_=*V*_0_=2.74 *µ*M) conditions and the corresponding results of the sensitivity analysis in **(g)** and **(h)** respectively for case 1 (both the inputs are absent). Normalized sensitivity matrix is shown with respect to the output (*P*). Here, yellow and blue correspond to the most sensitive and least sensitive values respectively. All the values of *P* were determined at 30 min. For cases 1 and 2, *R*_0_=0 *µ*M.

Similar to the OR gate, the XOR gate has two inputs, rapamycin and light, that restore the activity of *E* and *F* proteases respectively (Fig. 5c). However, to achieve an XOR functionality, we added additional reactions to the OR gate design in such a manner that these reactions can limit the output production only when both the inputs are present. In these reactions, protease *E* can degrade *U*, and protease *F* can degrade *A* and the intermediate complex *X*. Therefore, the two proteases can limit the production of each other, which limits the output production (Fig. 5c). For example, in the presence of rapamycin, the active protease *E* cleaves the substrate *S* to produce the output *P* and also degrades *U*. As in either of these reactions, *E* is not consumed, a high output should be produced. Similarly, in the presence of the light signal, a high output should be observed as the degradation of *A* (and *X*) by *F* does not affect the output activity (Fig. 5d). However, when both the inputs are present, *E* and *F* actively degrade *U* and *A* (and *X*) respectively, and this should limit the production of *P* to a minimal level.

Using an ODE model (see Supplementary Note S4), we determined the biosensor response for four different combinations of inputs (Fig. 5b). The same specification was used to categorize the output as either low or high as was used for the OR gate (*P*_1_>1 *µ*M and *P*_0_*<*0.5 *µ*M). Instead of observing a low output in case 4 (both the inputs are present), we observed a high output, that has almost the same activity as in case 2 (Fig. 5e). To achieve an XOR logical functionality for all input combinations, we then performed an optimization in *A*_0_, *B*_0_, *R*_0_, *U*_0_, and *V*_0_ analogously to what was done in the OR case and found a desired repose that met our specification (Fig. 5f).

To understand these results further, we conducted a sensitivity analysis for all four combinations of inputs. The results are shown for case 4 in Fig. 5g and h before and after optimizing the reaction conditions respectively. At the optimized condition, a reduced sensitivity of 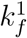 and 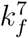 suggests reduction in the amount of *E* and *E*^′^ while at the same time, increased sensitivity of 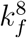 suggests an increased production of *F*. This leads to a higher sensitivity to the 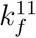 and 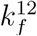 reaction parameters, which quantify how *E* and *F* degrade respectively *U* and *A* (and *X*), reducing the output in case 4. The results for the other cases are shown in Fig. S9. This type of analysis provides a way to understand the operation of a complex reaction network where several parameters govern the output dynamics.

## DISCUSSION

In the last decade, several biomolecular network topologies capable of performing computations similar to analog and digital circuits were proposed. These biomolecular circuits are based on gene expression, which requires multiple reactions to happen in a sequential order to produce an output, and because of that these circuits operate at a timescale of hours to days. This limits the computational complexity that can be achieved using molecular computation. Here, we presented a generalized framework to design a new class of protease-based Boolean logic gates for biosensing applications. These biosensors use post-translational protein modifications, which do not require a specific molecular queue, thereby allowing fast operation. We considered a realistic design framework to analyze the input-output characteristics of the CID and LID switches responsive to chemical and light input signals respectively. We then used these switches as elementary blocks to design biosensors to process two different inputs simultaneously, and produce either standard OR or XOR Boolean logic functionality. We improved the response of these biosensors through rigorous optimization, and the improvements were explained using sensitivity analysis. The biosensor’s capacity to meet the performance specifications considering biologically feasible reaction parameters suggests that this approach is viable for realistic chemical computing circuits.

The recent literature on designing of biomolecular logic gates lack a generalized approach that can be used to investigate the operation of an elementary circuit or a complex network.^29,30^ This is partially because of analyzing protease-based circuits can be difficult due to the irreversibility of cleavage reactions and a lack of a steady-state response. The mathematical optimization framework proposed here can be in principle adapted to model and understand the operation of a complex molecular circuit.

Some other designs and implementations of protein-based circuits have been proposed, that can mimic the operation of Boolean logic gates, but these designs cannot be tailored to biosensing applications.^8^ Moreover, most of the previously designed biosensing mechanisms can detect only one input at a time.^31–33^ Our approach is unique in the sense that we designed these sensors to recognize two different kinds of inputs simultaneously to produce a programmable output. However, the current design is limited in the sense that the biosensors cannot distinguish different sets of input combinations that provide the same output (either high or low). For example, the output of the OR gate is high for three possible scenarios: either or both the inputs are high. Similarly, for the XOR gate, we cannot distinguish between cases 2 and 3. To address this, a multiplexer-based approach might be used to design protease-based circuits where each input combination results in a unique output. ^34^

Cellular mechanisms use molecular computation to detect multiple chemical and physical input signals to execute an output that aids cellular function. Our approach can be extended to design other Boolean logic gates and eventually new multi-layered, multi-input circuits with complex network connectivity using enzymatic reactions that can be used to develop new types of biosensors with a capability of detecting multiple inputs.

## METHODS

The simulated response of each reaction network was determined by numerically integrating ODE models (shown in the Supplementary Materials) using the MATLAB ode23s solver unless otherwise specified. Initial conditions for each molecular species are described in figure captions, and the values of reaction parameters are shown in Table 1. To optimize the responses, we used the MATLAB fmincon function. We used the prerequisite conditions of each reaction as constraints to meet the specific performance criteria. For plots shown in Figs. 2e, 2f, 3e and 3f, to generate the 1000 combinations of kinetic parameters, each parameter was randomly sampled from a uniform distribution from an interval bounded by a lower bound of 0.1 × the nominal value and an upper bound of 10 × the nominal value given in Table 1. To determine the output sensitivity of each parameter, we calculated the sensitivity coefficient matrix over time (*s*_*i,j*_), which is defined as:^35^

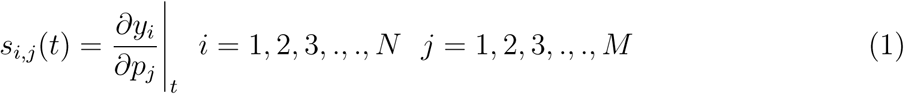

where *y*_*i*_ is a molecular species and *p*_*j*_ is a reaction parameter, the subscript *i* corresponds to a particular species, and the subscript *j* to a particular parameter in the system. In our study, *y*_*i*_ is *P* and *p*_*j*_ can be any of the parameters shown in Table 1. The ODE model equations can be written as:

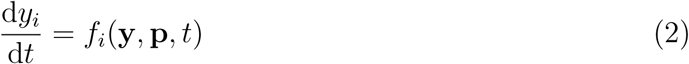

Here, **y** and **p** are the vectors of all the species and parameters, respectively. To calculate *s*_*i,j*_, we use a sensitivity differential equation, which can be expressed as:

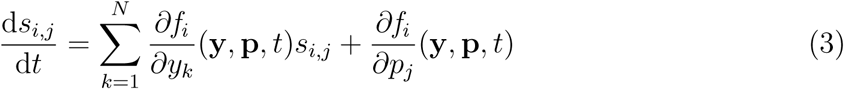

Equation (3) is solved numerically to calculate *s*_*i,j*_(*t*) for each parameter and the normalized values of 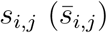 are reported in Figs. 4, 5, S8 and S9 using:

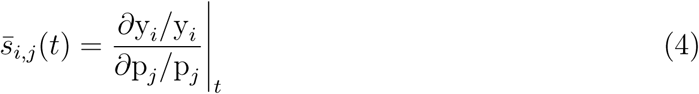

## Supporting information

Supp lnfo

## Acknowledgement

This research was supported in part by grants DARPA FA8650-18-1-7800 and NSF 1817936.

## Graphical TOC Entry

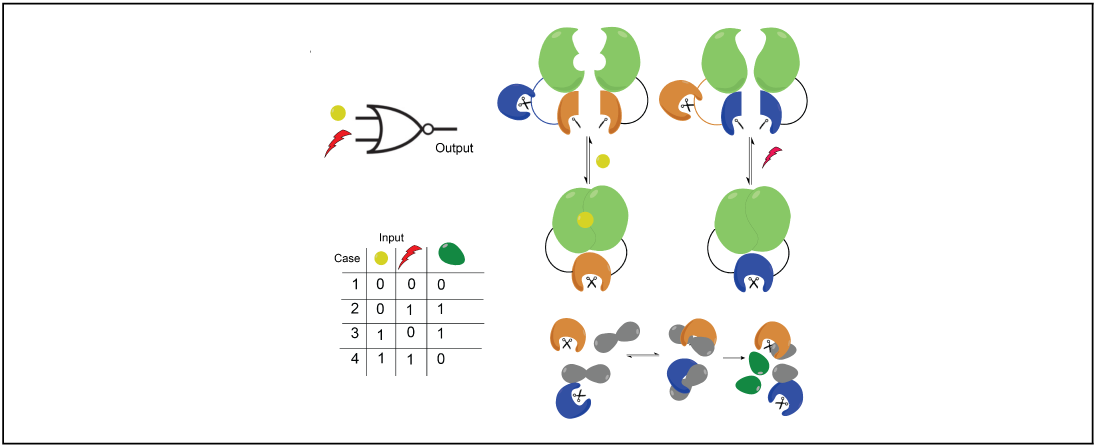

