## Supplementary material for "Mathematical models of protease-based enzymatic biosensors": Supp lnfo

### Supplemental Material
### Mathematical models of protease-based enzymatic biosensors

Deepak K. Agrawal,[†,‡] Sagar D. Khare,[*,¶] and Eduardo D Sontag[*,†,‡,§]

†*Department of Bioengineering, Northeastern University, Boston, M A, USA*

‡*Department of Electrical and Computer Engineering, Northeastern University, Boston, M A, USA*

¶*Department of Chemistry and Chemical Biology, Rutgers University, Piscataway, NJ, USA*

§*Laboratory of Systems Pharmacology, Program in Therapeutic Science, Harvard Medical School, Boston, M A, USA*

### Supplementary Note S1

Using the mass-action law, the kinetics of the CID switch can be modeled as:

$$\dot{A} = k_r^1 X - k_f^1 AR + k_r^4 E - k_f^4 YA - k_f^7 AB + k_r^7 E', \tag{1}$$

$$\dot{R} = k_r^1 X - k_f^1 AR + k_r^2 Y - k_f^2 BR, \tag{2}$$

$$\dot{B} = k_r^2 Y - k_f^2 BR + k_r^3 E - k_f^3 XB - k_f^7 AB + k_r^7 E', \tag{3}$$

$$\dot{X} = k_f^1 AR - k_r^1 X - k_f^3 XB + k_r^3 E, \tag{4}$$

$$\dot{Y} = k_f^2 BR - k_r^2 Y - k_f^4 YA + k_r^4 E, \tag{5}$$

$$\dot{E} = k_f^4 YA + k_f^3 XB - k_r^3 E - k_r^4 E - k_f^5 ES + k_r^5 C + k_f^6 C, \tag{6}$$

$$\dot{S} = -k_f^5 ES + k_r^5 C - k_f^5 E'S + k_r^5 C', \tag{7}$$

$$\dot{C} = k_f^5 ES - k_r^5 C - k_f^6 C, \tag{8}$$

$$\dot{P} = k_f^6 C + k_f^6 C', \tag{9}$$

$$\dot{E'} = k_f^7 AB - k_r^7 E' - k_f^5 E'S + k_r^5 C' + k_f^6 C', \tag{10}$$

$$\dot{C'} = k_f^5 E'S - k_r^5 C' - k_f^6 C'. \tag{11}$$

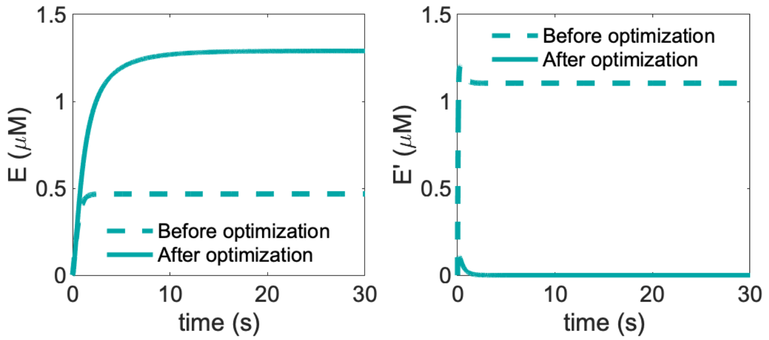


Figure S1: Timing response of the CID switch in the presence of input ($R$) before optimization where $A_0$=$B_0$=5 $\mu$M, and $R_0$=0.5 $\mu$M (dashed-line), and after optimization where $A_0$=$B_0$=1.44 $\mu$M and $R_0$=3.01 $\mu$M (solid-line). A reduction in the initial concentrations of $A$ and $B$ reduced the amount of $E$ and $E'$, but an increase in $R_0$ increased only the amount of $E$, resulting in an overall increase in the amount of $E$. The ODE model shown in Note S1 was used to simulate the response with parameters shown in Table 1. Here $S_0$ was 5 $\mu$M while the rest of the molecular species were initially set to 0 $\mu$M.

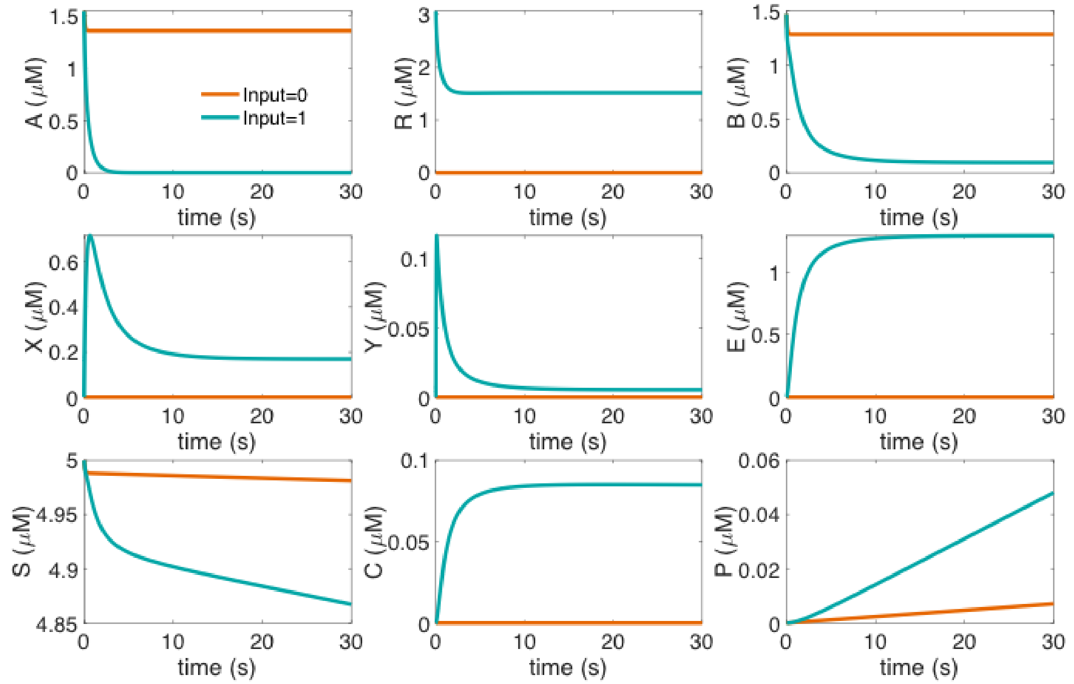


Figure S2: Simulated timing response of different species that make up the CID switch while keeping $A_0$=$B_0$=5 $\mu$M, and $R_0$=0.5 $\mu$M. Species involved in the sensing element reached steady-state within a few seconds. The ODE model shown in Note S1 was used to simulate the response with parameters shown in Table 1. Here $S_0$ was 5 $\mu$M while the rest of the molecular species were initially set to 0 $\mu$M.

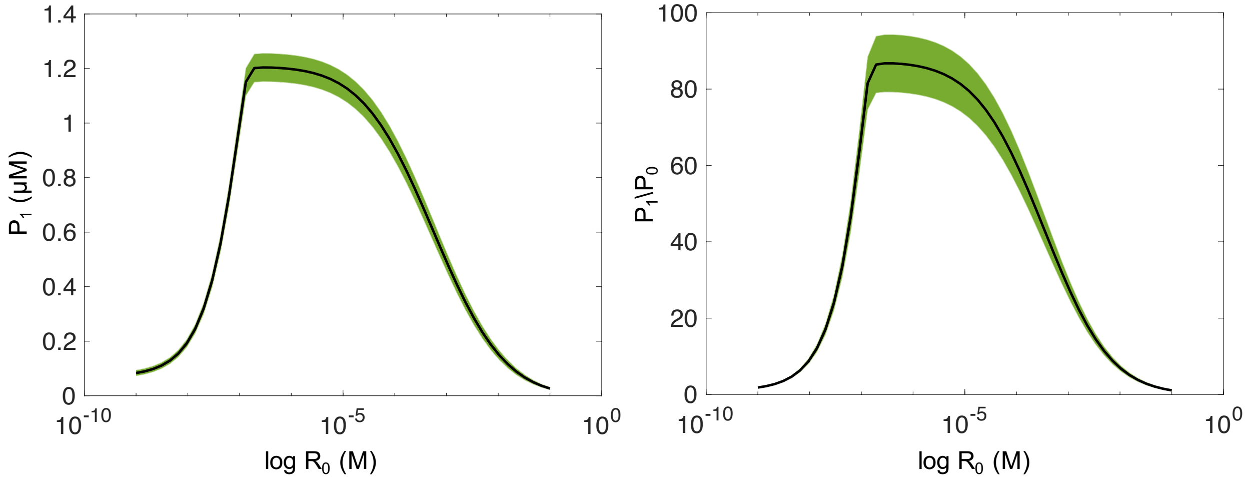


Figure S3: Performance evaluation of the CID switch as a function of $R_0$ while keeping $A_0$ and $B_0$ fixed at 0.144 $\mu$M each. For each concentration of $R_0$, 1,000 simulations were conducted. Parameter values were randomly sampled from a uniform distribution (see Methods). The ODE model shown in Note S1 was used to simulate the response with parameters shown in Table 1. Averaged metrics of $P_1$ and the $P_1/P_0$ are shown here. All the values of $P$ were determined at 30 min. The error bars are shown in the shaded region and were determined using the standard error of the mean. Here $S_0$ was 5 $\mu$M while the rest of the molecular species were initially set to 0 $\mu$M.

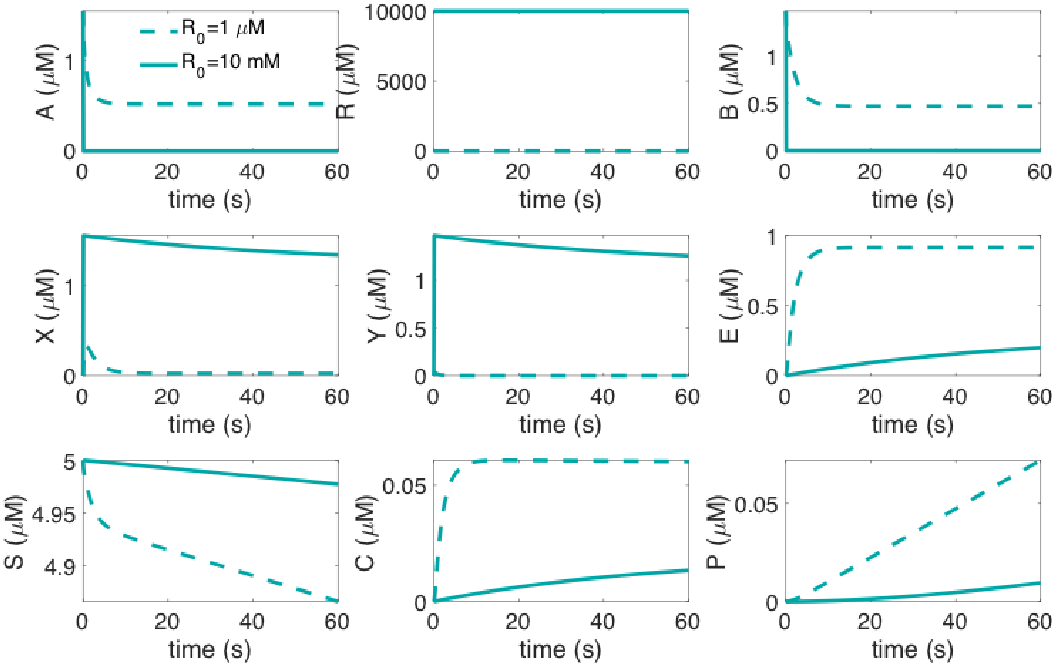


Figure S4: Simulated timing response of different species that make up the CID switch at different initial concentrations of $R$ while keeping $A_0$ and $B_0$ fixed at 1.44 $\mu$M each. An increase in $R_0$, led to a higher interaction between $B$ and $R$, and because of that, less $B$ is available to bind to $X$ in order to form $E$. The ODE model shown in Note S1 was used to simulate the response with parameters shown in Table 1. Here $S_0$ was 5 $\mu$M while the rest of the molecular species were initially set to 0 $\mu$M.

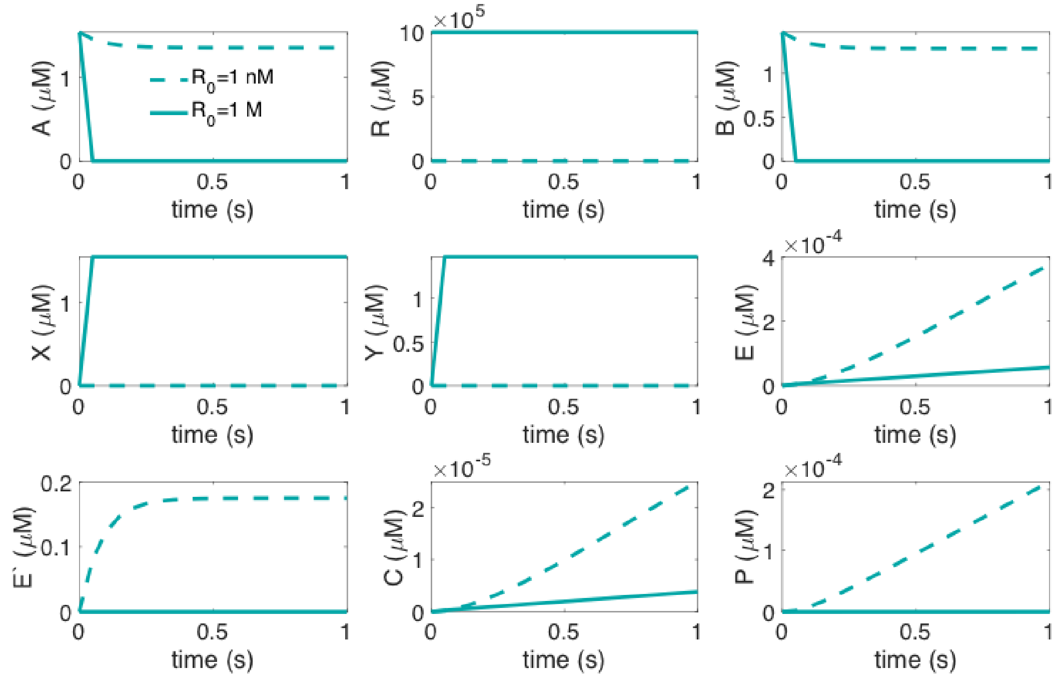


Figure S5: Simulated timing response of different species that make up the CID switch at different initial concentrations of $R$ while keeping $A_0$ and $B_0$ fixed at 1.44 $\mu$M each. At the lower concentrations of $R_0$, $A$ and $B$ are freely available to produce $E'$ than at the higher values of $R_0$ where $A$ and $B$ are sequestered by $R$, and forms $X$ and $Y$. The ODE model shown in Note S1 was used to simulate the response with parameters shown in Table 1. Here $S_0$ was 5 $\mu$M while the rest of the molecular species were initially set to 0 $\mu$M.

### Supplementary Note S2

Using the mass-action law, the kinetics of the LID switch can be modeled as:

$$\dot{U} = -k_f^8 UV + k_r^8 F, \text{(dark)} \tag{12}$$

$$\dot{V} = -k_f^8 UV + k_r^8 F, \text{(dark)} \tag{13}$$

$$\dot{U} = -k_f^8 UV + k_r^9 F, \text{(light)} \tag{14}$$

$$\dot{V} = -k_f^8 UV + k_r^9 F, \text{(light)} \tag{15}$$

$$\dot{F} = k_r^5 D + k_f^6 D - k_f^5 FS + k_f^8 UV - k_r^8 F, \tag{16}$$

$$\dot{S} = -k_f^5 FS + k_r^5 D, \tag{17}$$

$$\dot{D} = k_f^5 FS - k_r^5 D - k_f^6 D, \tag{18}$$

$$\dot{P} = k_f^6 D. \tag{19}$$

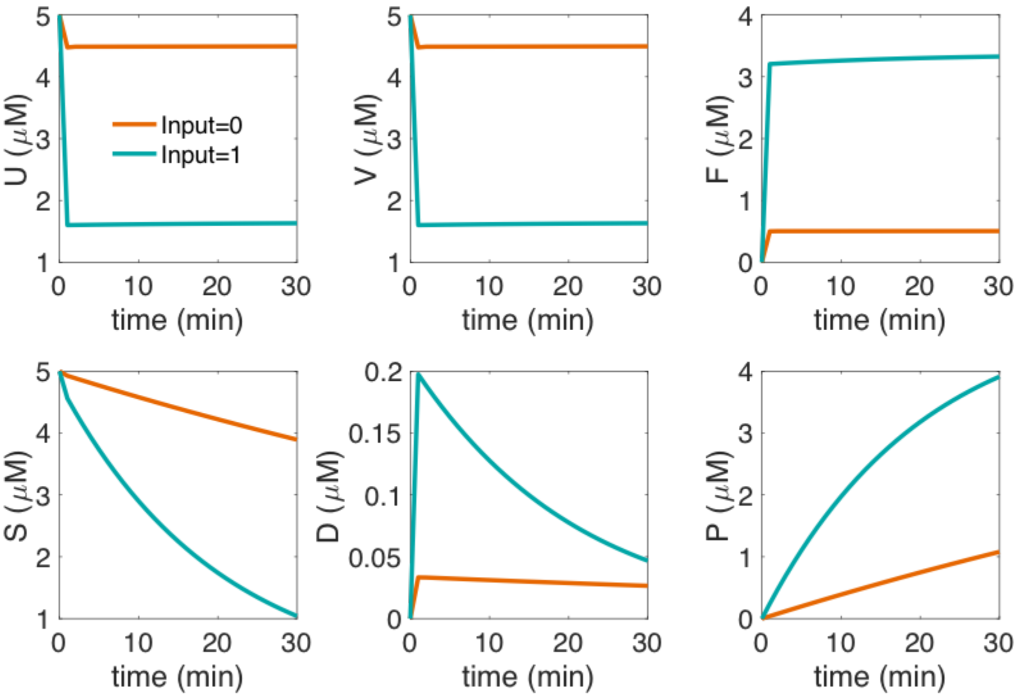


Figure S6: Simulated timing response of different species that make up the LID switch in the absence (denoted as 0) and presence of input (denoted as 1) while keeping $U_0$ and $V_0$ at 5 $\mu$M each. $U$ and $V$ dimerized even in the absence of input, which allowed to form $F$, resulted in a high amount of $P$. The ODE model shown in Note S2 was used to simulate the response with parameters shown in Table 1. Here $S_0$ was 5 $\mu$M while the rest of the molecular species were initially set to 0 $\mu$M.

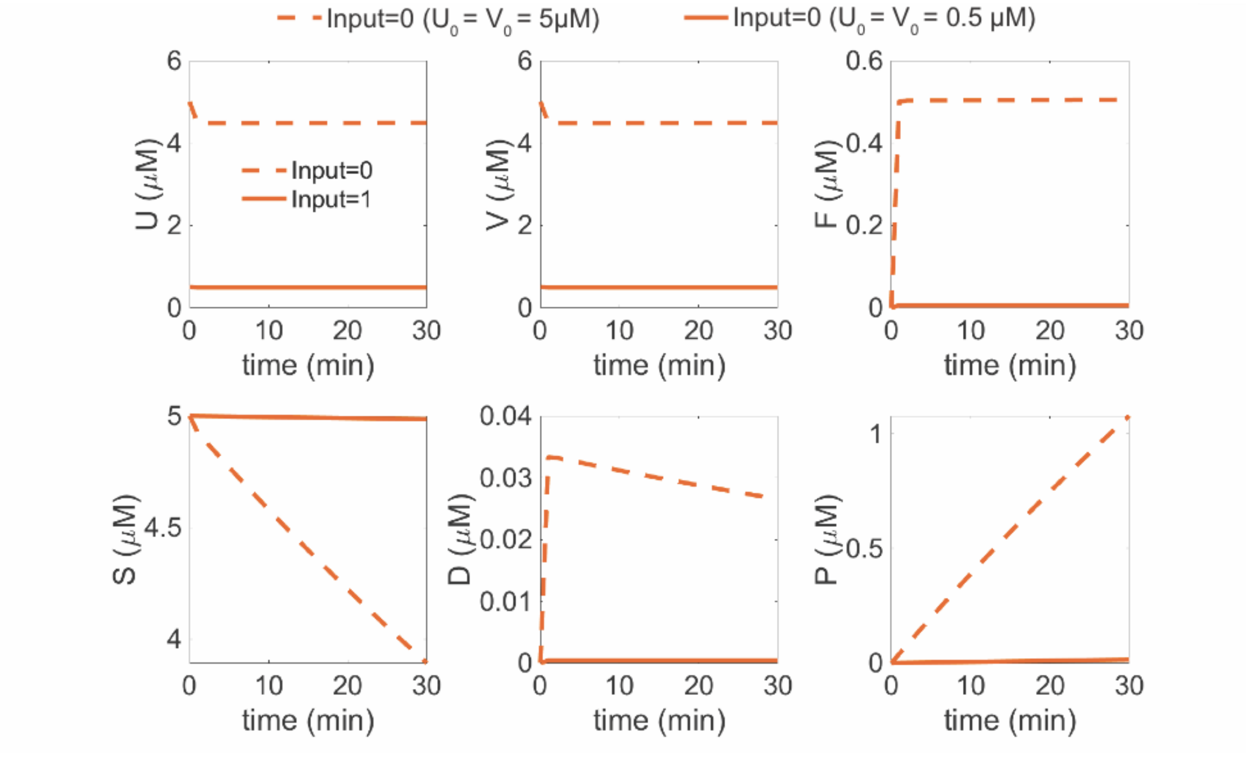


Figure S7: Simulated timing response of different species that make up the LID switch in the absence of input while keeping $U_0$ and $V_0$ fixed at 5 $\mu$M (dashed-line) or at 0.5 $\mu$M (solid-line) each. At a reduced initial concentrations of $U$ and $V$, $F$ is almost negligible. The ODE model shown in Note S2 was used to simulate the response with parameters shown in Table 1. Here $S_0$ was 5 $\mu$M while the rest of the molecular species were initially set to 0 $\mu$M.

### Supplementary Note S3

Using the mass-action law, the kinetics of the OR gate can be modeled as:

$$\dot{A} = k_r^1 X - k_f^1 AR + k_r^4 E - k_f^4 YA - k_f^7 AB + k_r^7 E', \tag{20}$$

$$\dot{R} = k_r^1 X - k_f^1 AR + k_r^2 Y - k_f^2 BR, \tag{21}$$

$$\dot{B} = k_r^2 Y - k_f^2 BR + k_r^3 E - k_f^3 XB - k_f^7 AB + k_r^7 E', \tag{22}$$

$$\dot{X} = k_f^1 AR - k_r^1 X - k_f^3 XB + k_r^3 E - k_f^{12} FX + k_r^{12} I, \tag{23}$$

$$\dot{Y} = k_f^2 BR - k_r^2 Y - k_f^4 YA + k_r^4 E, \tag{24}$$

$$\dot{E} = k_f^4 YA + k_f^3 XB - k_r^3 E - k_r^4 E - k_f^5 ES + k_r^5 C + k_f^6 C, \tag{25}$$

$$\dot{S} = -k_f^5 ES + k_r^5 C - k_f^5 FS + k_r^5 D - k_f^5 E'S + k_r^5 C', \tag{26}$$

$$\dot{C} = k_f^5 ES - k_r^5 C - k_f^6 C, \tag{27}$$

$$\dot{U} = -k_f^8 UV + k_r^8 F, \text{(dark)} \tag{28}$$

$$\dot{V} = -k_f^8 UV + k_r^8 F, \text{(dark)} \tag{29}$$

$$\dot{U} = -k_f^8 UV + k_r^9 F, \text{(light)} \tag{30}$$

$$\dot{V} = -k_f^8 UV + k_r^9 F, \text{(light)} \tag{31}$$

$$\dot{F} = k_r^5 D + k_f^6 D - k_f^5 FS + k_f^8 UV - k_r^8 F \tag{32}$$

$$- k_f^{12} FX + k_r^{12} I + k_f^{13} I, \tag{33}$$

$$\dot{D} = k_f^5 FS - k_r^5 D - k_f^6 D, \tag{34}$$

$$\dot{P} = k_f^6 C + k_f^6 D + k_f^6 C', \tag{35}$$

$$\dot{E}' = k_f^7 AB - k_r^7 E' - k_f^5 E'S + k_r^5 C' + k_f^6 C', \tag{36}$$

$$\dot{C}' = k_f^5 E'S - k_r^5 C' - k_f^6 C'. \tag{37}$$

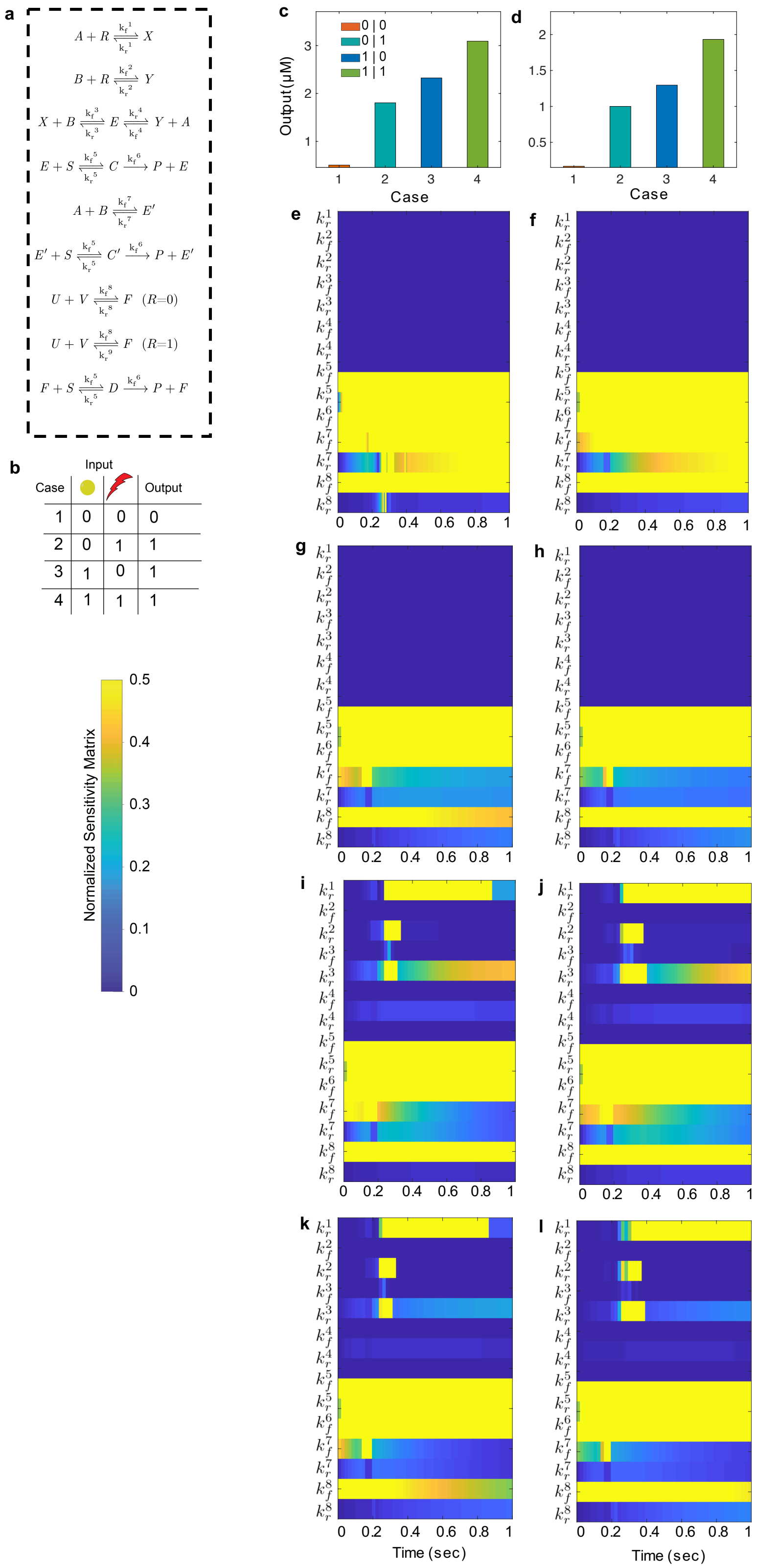


Figure S8: Protease based Boolean OR gate. (a) Chemical reactions of the OR gate and (b) the corresponding truth table. The ODE model shown in Note S3 was used to simulate the response with parameters shown in Table 1. (c-d) Simulated response of the OR gate at (c) unoptimized ($A_0$=$B_0$=1.44 $\mu$M, $R_0$=3.01 $\mu$M, and $U_0$=$V_0$=1.59 $\mu$M) and **(f)** optimized ($A_0$=$B_0$=0.72 $\mu$M, $R_0$=1.67 $\mu$M, and $U_0$=$V_0$=1 $\mu$M) conditions and the corresponding results of the sensitivity analysis are shown in (e) and (f) for case 1; in (g) and (h) for case 2; in (i) and (j) for case 3; in (k) and (l) for case 4, respectively. Normalized sensitivity matrix is shown with respect to the output ($P$). Here, yellow and blue correspond to the most sensitive and least sensitive values respectively. All the values of $P$ were determined at 30 min. For cases 1 and 2, $R_0$=0 $\mu$M while in cases 2 and 4, $k_r^8$ was replaced with $k_r^9$. Here, $S_0$ was 5 $\mu$M while the rest of the molecular species were initially set to 0 $\mu$M.

### Supplementary Note S4

Using the mass-action law, the kinetics of the XOR gate can be modeled as:

$$\dot{A} = k_r^1 X - k_f^1 AR + k_r^4 E - k_f^4 YA - k_f^7 AB + k_r^7 E' - k_f^{12} FA + k_r^{12} H, \tag{38}$$

$$\dot{R} = k_r^1 X - k_f^1 AR + k_r^2 Y - k_f^2 BR, \tag{39}$$

$$\dot{B} = k_r^2 Y - k_f^2 BR + k_r^3 E - k_f^3 XB - k_f^7 AB + k_r^7 E', \tag{40}$$

$$\dot{X} = k_f^1 AR - k_r^1 X - k_f^3 XB + k_r^3 E - k_f^{12} FX + k_r^{12} I, \tag{41}$$

$$\dot{Y} = k_f^2 BR - k_r^2 Y - k_f^4 YA + k_r^4 E, \tag{42}$$

$$\dot{E} = k_f^4 YA + k_f^3 XB - k_r^3 E - k_r^4 E - k_f^5 ES + k_r^5 C + k_f^6 C \tag{43}$$

$$- k_f^{10} EU + k_r^{10} G + k_f^{11} G, \tag{44}$$

$$\dot{S} = -k_f^5 ES + k_r^5 C - k_f^5 FS + k_r^5 D - k_f^5 E'S + k_r^5 C', \tag{45}$$

$$\dot{C} = k_f^5 ES - k_r^5 C - k_f^6 C, \tag{46}$$

$$\dot{U} = -k_f^8 UV + k_r^8 F - k_f^{10} EU + k_r^{10} G, \text{(dark)} \tag{47}$$

$$\dot{V} = -k_f^8 UV + k_r^8 F, \text{(dark)} \tag{48}$$

$$\dot{U} = -k_f^8 UV + k_r^9 F - k_f^{10} EU + k_r^{10} G, \text{(light)} \tag{49}$$

$$\dot{V} = -k_f^8 UV + k_r^9 F, \text{(light)} \tag{50}$$

$$\dot{F} = k_r^5 D + k_f^6 D - k_f^5 FS + k_f^8 UV - k_r^8 F - k_f^{12} FA + k_r^{12} H + k_f^{13} H \tag{51}$$

$$- k_f^{12} FX + k_r^{12} I + k_f^{13} I, \tag{52}$$

$$\dot{D} = k_f^5 FS - k_r^5 D - k_f^6 D, \tag{53}$$

$$\dot{P} = k_f^6 C + k_f^6 D + k_f^6 C', \tag{54}$$

$$\dot{E'} = k_f^7 AB - k_r^7 E' - k_f^5 E'S + k_r^5 C' + k_f^6 C', \tag{55}$$

$$\dot{C'} = k_f^5 E'S - k_r^5 C' - k_f^6 C', \tag{56}$$

$$\dot{G} = k_f^{10} EU - k_r^{10} G - k_f^{11} G, \tag{57}$$

$$\dot{U}' = k_f^{11} G, \tag{58}$$

$$\dot{H} = k_f^{12} F A - k_r^{12} H - k_f^{13} H, \tag{59}$$

$$\dot{I} = k_f^{12} F X - k_r^{12} I - k_f^{13} I, \tag{60}$$

$$\dot{A}' = k_f^{13} H, \tag{61}$$

$$\dot{X}' = k_f^{13} I. \tag{62}$$

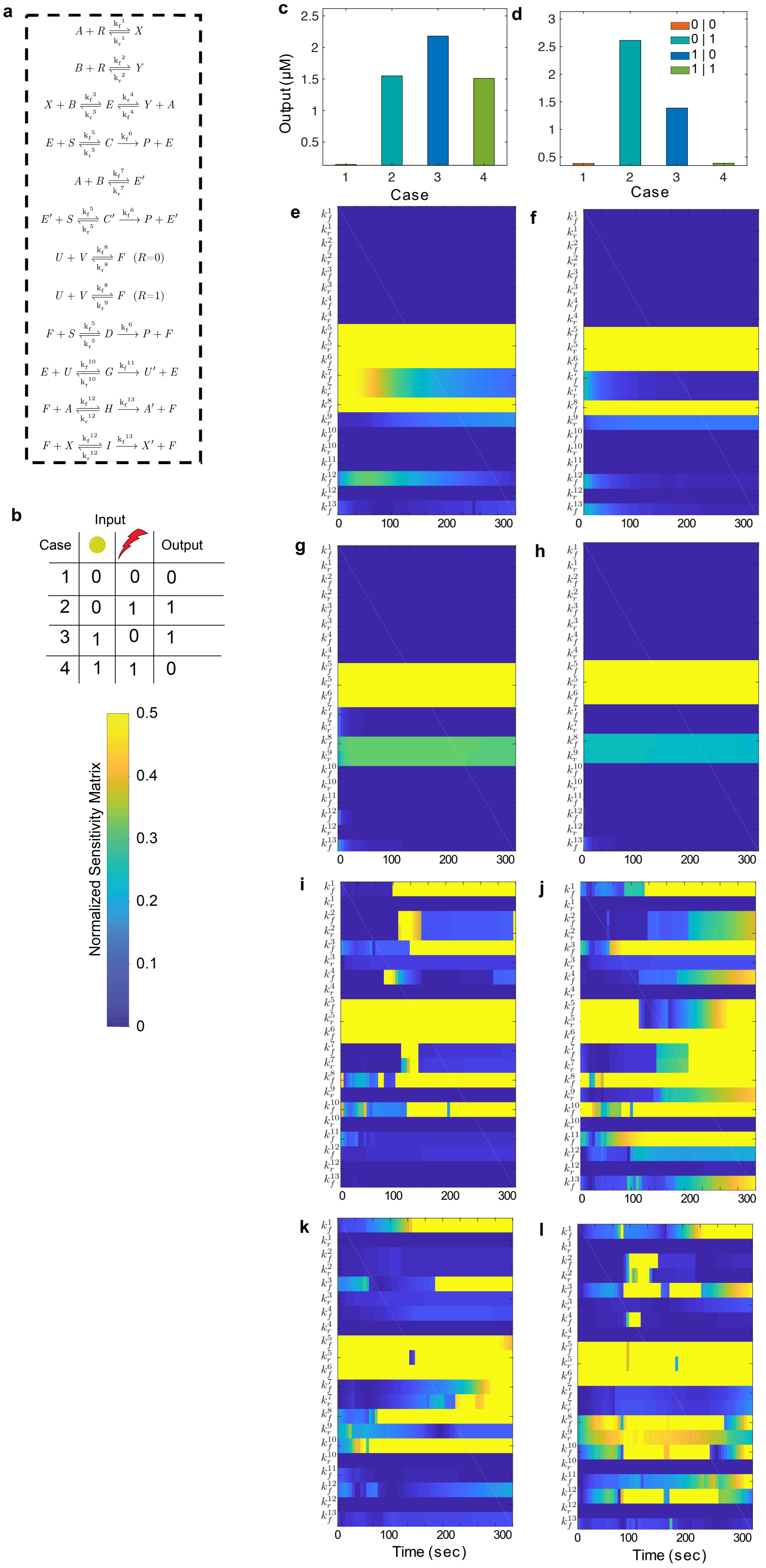


Figure S9: Protease based Boolean XOR gate. (a) Chemical reactions of the XOR gate and (b) the corresponding truth table. The ODE model shown in Note S4 was used to simulate the response with parameters shown in Table 1. (c-d) Simulated response of the OR gate at (c) unoptimized ($A_0$=$B_0$=1.44 $\mu$M, $R_0$=3.01 $\mu$M, and $U_0$=$V_0$=1.59 $\mu$M) and **(f)** optimized ($A_0$=1.22 $\mu$M, $B_0$=0.88 $\mu$M, $R_0$=0.94 $\mu$M, and $U_0$=$V_0$=2.74 $\mu$M) conditions and the corresponding results of the sensitivity analysis are shown in (e) and (f) for case 1; in (g) and (h) for case 2; in (i) and (j) for case 3; in (k) and (l) for case 4, respectively. Normalized sensitivity matrix is shown with respect to the output ($P$). Here, yellow and blue correspond to the most sensitive and least sensitive values respectively. All the values of $P$ were determined at 30 min. For cases 1 and 2, $R_0$=0 $\mu$M while in cases 2 and 4, $k_r^8$ was replaced with $k_r^9$. Here, $S_0$ was 5 $\mu$M while the rest of the molecular species were initially set to 0 $\mu$M.
